# Visual information asymmetry shapes leadership, gaze, and interaction control strategies in joint object manipulation

**DOI:** 10.64898/2026.11.18.767522

**Authors:** Alireza Khaleghi, Feifan Huang, Dorian Verdel, Etienne Burdet

## Abstract

When two people carry a large object, they often have different visual information about the surrounding environment. For example, when carrying a table through a doorway, the person at the front can see the path ahead, whereas the person at the back can monitor nearby obstacles to avoid collisions. How such sensory asymmetry affects coordination, and whether haptic feedback improves performance, remain unclear. We investigated these questions in a co-manipulation task involving 12 dyads. Each partner controlled one end of a virtual canoe through a wrist robot while navigating a winding river. We varied visual view, with the rear partner able or not able to see the path ahead of the canoe, and independently provided haptic feedback to neither, one, or both partners. All dyads successfully completed the task under both visual conditions. When only the front partner could see the path ahead, conditional Granger causality revealed stronger influence from this partner, whereas influence was balanced when both partners could see ahead. When compared to symmetric visual information, the asymmetric condition increased the canoe’s lateral position and reduced the angular deviation relative to the river centerline heading. The front partner also made smaller and more frequent saccades, and adapted the effective observation space. Haptic feedback did not substantially improve overall task performance. A simple model in which each partner independently corrected their end of the canoe using delayed visual feedback, without relying on an explicit model of the object or the partner’s actions, reproduced the main behavioral differences between the asymmetric and symmetric visual conditions. These findings suggest that unequal visual information shapes coordination roles, movement patterns, and eye gaze behavior during physical collaboration.

## I. Introduction

**P**eople routinely and seamlessly manipulate large objects together in constrained environments, such as when carrying a table together through a narrow doorway. In such scenarios the object or the environment can restrict the partners’ visual information and constrain their movements. For instance, the person at the front may be able to see the path ahead, whereas the person at the back must monitor whether the table is colliding with a wall. The partners therefore have complementary visual information that they may exchange through haptic interactions [1]. In the present paper, we systematically vary the visual and haptic information available to each partner to investigate the respective roles of these sensory modalities in physical cooperation and the strategies that partners use to coordinate their actions.

Previous studies have shown that people coordinate their actions in space and time by sharing task representations and predicting each other’s actions [1]–[3] Physical coupling generally improves motor performance [4]–3] and interpersonal coordination [8]. Many studies have described such interactions in terms of leader-follower roles [4], [8], [9]. However, definitions of these roles vary across studies, and apparent leadership may arise from task constraints rather than stable differences between partners [10]. Task constraints and differences in available information may therefore influence the emergence of coordination roles.

In tasks involving haptic feedback, incomplete information about a partner can alter the strategy used for collaboration [11]. Removing visual feedback to one partner can reduce joint performance and change interpersonal coordination [12], [13]. Differences between partners can arise from unequal mechanical capabilities [14], [15] or from differences in the quality of the information available to them: When physically connected partners have unequal visual or haptic information about a target, the partner with more complete or more accurate information tends to drive the interaction [16]–[18].

Joint object carrying has also been studied more directly. In [19], dyads of participants carried a table to nine goal locations, with either one or both partners knowing the goal. Their movement strategies varied according to whether one or both partners had this information. However, the study did not examine task-space constraints imposed by obstacles or degraded visual information. [18] extended this work by having participants carry a virtual object through an environment comprising several rooms. That study focused on decision-making and did not consider the case of asymmetric visual information with respect to upcoming obstacle and paths arising from first-person viewpoints. In [20], leader and follower roles were assigned in advance rather than allowed to emerge from the partners’ access to information about the environment.

We designed an experiment that captures key constraints of object co-manipulation while allowing visual and haptic information to be controlled independently. We selected a task that reproduced the constraints of carrying a table through a doorway while requiring continuous sensory feedback. Each of two partners controlled one end of a virtual canoe using a wrist robotic interface, and jointly steered it along a winding river (Fig 1a). We manipulated (i) the rear partner’s available visual information to reproduce the occlusion encountered during object carrying, and (ii) the haptic feedback provided to each partner in each visual information scenario to examine its contribution to coordination and performance. We tested four hypotheses:

H1: Despite having limited but complementary visual information, partners develop a successful co-manipulation strategy.

H2: Access to task-relevant information affects both partners’ control strategies and visual behavior.

H3: Consistent with the emergence of leader–follower roles under unequal information access, the partner with more useful visual information takes the lead in the dyad.

H4: Haptic feedback about interactions with the shared object improves performance in the presented collaborative task.

**Fig. 1:**
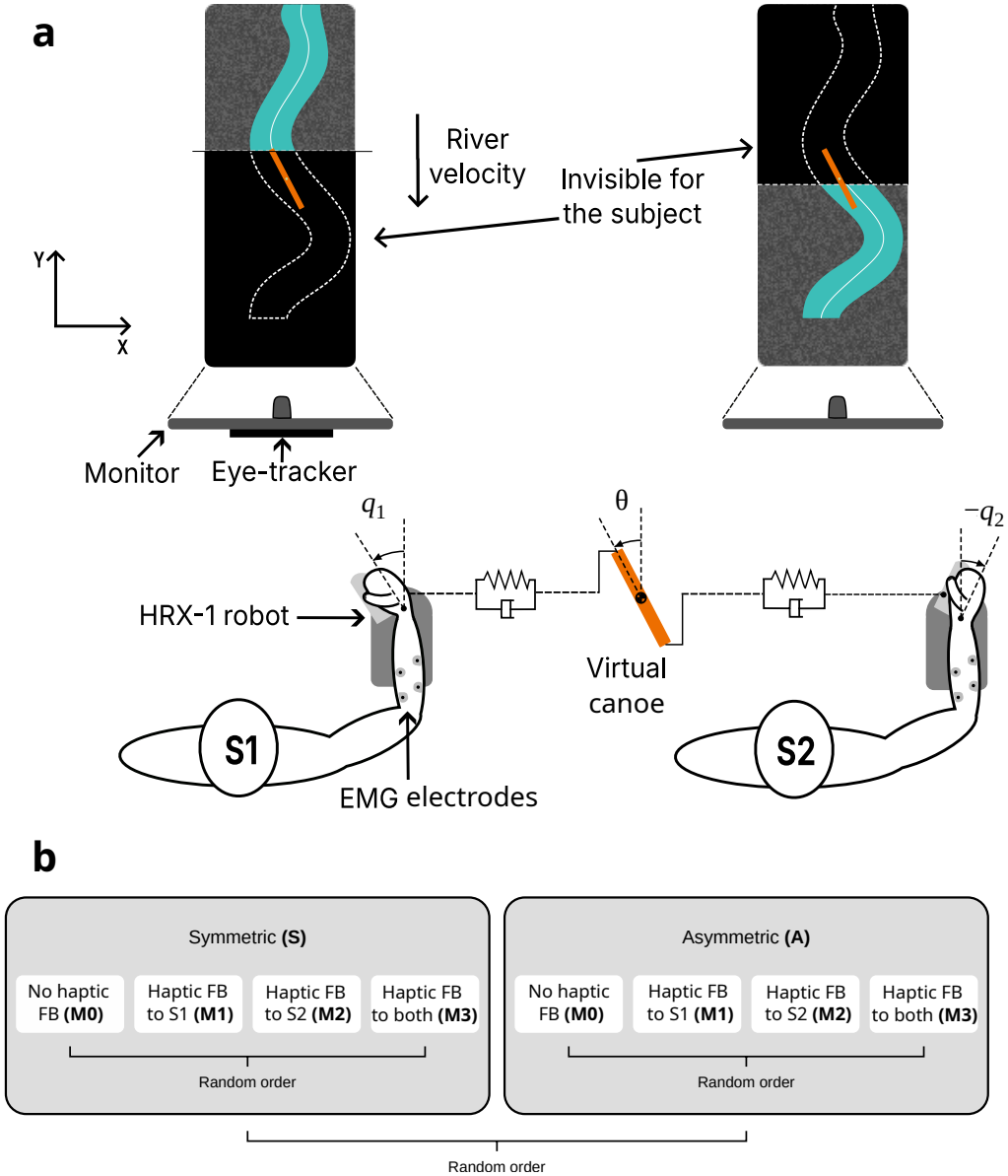
Experiment’s overview. **(a)** Two participants controlled a virtual canoe through wrist robots and spring-damper couplings. **(b)** Protocol with the two visual view conditions (Symmetric, Asymmetric) and four haptic feedback modes (M0–M3).

## II. Methods

### A. Experiment

The study was approved by the Science, Engineering and Technology Research Ethics Committee of Imperial College London (SETREC reference 7112072). Twenty-four participants (8 female and 16 male, aged 21–42 years) took part in the study as 12 dyads. Participants were recruited from students and staff of Imperial College. Inclusion criteria were right-handedness, normal or corrected-to-normal vision, and no self-reported neurological or musculoskeletal condition affecting the wrist. All participants provided written informed consent before the experiment.

Each dyad navigated a virtual canoe along a river (Fig. 1a). The participants sat side by side, separated by an opaque curtain, and viewed the virtual task on an individual monitor. The monitors were identical and measured 60 × 34 cm, which corresponds to 924 × 520 l.u. (length unit), which is used throughout the rest of the paper for reporting length measures of the virtual task. Participants were instructed not to communicate verbally during the task. Each participant controlled a cursor through wrist flexion and extension using an HRX-1 wrist robotic interface (Human RobotiX, London, UK). The two cursors were connected to the canoe’s bow and stern via viscoelastic couplings. The dyad was instructed to keep the canoe within the river.

The virtual canoe was modeled as a rigid bar of length *L* = 150 l.u. × *W* = 15 l.u., with a virtual mass *M* = 5 kg, and moment of inertia *J* = 4000 Nm.s^2^/rad. Its center was constrained to a constant vertical position, so that its configuration is described by its lateral position and orientation (*x*_*c*_, *θ*). Let *c*_*i*_ denote the lateral position of the cursor controlled by participant *S*_*i*_ (*i* = 1, 2), and *x*_*i*_ denote the corresponding lateral position of the canoe attachment point:

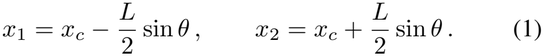

The force exerted on the canoe through each viscoelastic coupling was then calculated as

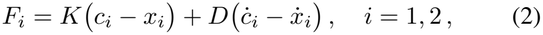

where *K* = 15 N/l.u. and *D* = 0.05 Nm.s/rad denote the coupling stiffness and damping, respectively. Using Eq. 2, Newton’s second law yields

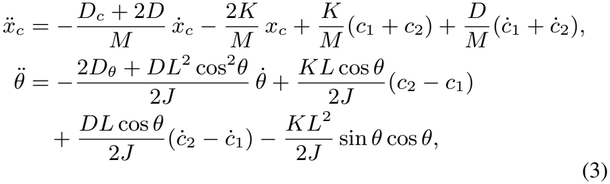

where *D*_*c*_ = 20 N.s/l.u. and *D*_*θ*_ = 4000 Nm.s/rad denote the translational and rotational viscous damping of the canoe moving in the river.

The canoe’s two degrees of freedom *x*_*c*_ and *θ* were controlled jointly by the two participants, but neither participant could control either degree of freedom independently. If the canoe crossed a river bank, the river stopped and both participants received a 0.5 s vibration at the wrist. The task was implemented in Unity 2022.3.18f1, with haptic control implemented in Simulink (MATLAB R2023b). A cursor displacement of 250 l.u. corresponded to 1 rad of wrist rotation, and the river scrolled at 48 l.u./s.

The experiment used two visual view conditions and four haptic feedback modes. Under symmetric view (S), both participants could see the upcoming river. Under asymmetric view (A), S1 could see only the upcoming river, from the vertical position of their cursor to the top of the monitor, whereas S2 could see only the section already passed, from the canoe’s center to the bottom of the monitor (Fig. 1a). The canoe and both cursors remained visible to both participants in all conditions. The spring-damper coupling of Eq. 2 generated an interaction force at each end of the canoe. The corresponding wrist torque was fed back to: neither subject (M0), S1 only (M1), S2 only (M2), or to both subjects (M3). Canoe dynamics and cursor couplings were identical across feedback modes. Boundary vibrations were provided in all conditions, including M0.

Each participant *i* ∈ {1, 2} received haptic feedback of the interaction with the canoe in the form of virtual force *F*_*i*_ rendered as a wrist torque 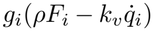, where *ρ* = 5 · 10^*−*4^ Nm/l.u. converts the force to torque and *k*_*v*_ = 0.1 Nm.s/rad is a viscous resistance added to avoid chattering. *g*_*i*_ ∈ {0, 1} encodes the haptic mode: *g*_1_ = *g*_2_ = 0 in M0, *g*_1_ = 1, *g*_2_ = 0 in M1, *g*_1_ = 0, *g*_2_ = 1 in M2, and *g*_1_ = *g*_2_ = 1 in M3. The main task parameters and definitions are listed in Table I.

**TABLE I:** List of experimental parameters.

|  | **Definition** | **Value** |
|---|---|---|
| *S*<sub>1</sub> | Front partner | - |
| *S*<sub>2</sub> | Rear partner | - |
| *x*<sub>*c*</sub> | Lateral position of canoe center | - |
| *θ* | Heading of the virtual canoe | - |
| *L* | Length of the canoe | 150 l.u. |
| *W* | Width of the canoe | 15 l.u. |
| *M* | Mass of the canoe | 5 kg |
| *J* | Inertia of the canoe | 4000 Nm.s<sup>2</sup>/rad |
| *W*<sub>*r*</sub> | Width of the river | 80 l.u. |
| *v* | Descend speed of the river | 48 l.u./s |
| △ | Lead/lag of subjects w.r.t. canoe center | 1.56 s |
| *c*<sub>1</sub>, *c*<sub>2</sub> | Lateral positions of participant's cursors | - |
| *x*<sub>1</sub>, *x*<sub>2</sub> | Lateral positions of tips of the canoe | - |
| *q*<sub>1</sub>, *q*<sub>2</sub> | Angle of robotic wrist interface | - |
| *F*<sub>1</sub>, *F*<sub>2</sub> | Virtual connection forces | - |
| *x*<sub>1</sub>, *x*<sub>2</sub> | Lateral positions of tips of the canoe | - |
| *g*<sub>*i*</sub> | Haptic feedback switch | 0 or 1 |
| *s* | Scale factor of JZS prior | - |
| BF | Bayes factor | - |
| *d*<sub>*z*</sub> | Cohen's effect size | - |

Each dyad first completed a familiarization trial with symmetric view and bilateral haptic feedback. The experiment then comprised eight blocks of five trials, with the blocks covering all combinations of view condition and feedback mode (Fig. 1b). The order of view conditions was randomized, and the order of haptic feedback modes was randomized within each view condition. Seven dyads started with the asymmetric view and five with the symmetric view. The same five river profiles were presented in a fixed order within each block. Each trial included 69.0 ± 7.0 *s* of river motion, where the deviation stems from pauses caused by hitting the edge.

### B. Measurement and Analysis

Muscle activity was recorded but is not analyzed in the Results, as it was found to mainly reflect the mechanical load associated with task execution without clear effects of visual or haptic information availability.

#### 1) Gaze

The gaze position of S1, the front participant, was recorded at 120 Hz using a Tobii Pro Fusion eye tracker. The eye tracker was calibrated before each view condition, and gaze coordinates were transformed into the task reference frame. Gaze was taken from the left eye. Samples falling outside the screen were discarded, and gaps shorter than 75 ms were linearly interpolated. Gaze positions were converted to degrees of visual angle using the screen dimensions (60 × 34 cm) and a viewing distance of 60 cm, which was fixed through a head movement stabilizer. Trials with less than 5 s of valid tracking were excluded. Saccades were identified using the velocity-threshold of 30^*°*^ /s [21]. Dyad 11 was removed due a bias in their gaze data. Saccade frequency was defined as the number of saccades divided by the valid tracking time. Saccade amplitude and peak velocity were averaged over the saccades of each trial. The mean gaze position was computed by averaging the gaze data along the x and y axes over the trial duration.

#### 2) Robot States

Robot and task states were recorded at 74.4 ± 5.4 Hz (mean ± SD across trials). Each trial was analyzed from the moment the canoe entered the river (3.9 ± 2.6 s after the start of the recording, mean ± SD across trials) until it exited the river (69.0 ± 7.0 s). Recorded signals included wrist angles, angular velocities, commanded torques, cursor positions, interaction forces, canoe kinematics, and river geometry. Simulation-derived states and interaction forces were analyzed without additional filtering. Processed gaze signals were linearly interpolated onto the robot timestamps. Of the 480 planned trials, two were not recorded and two were incomplete (less than 55 s of river motion); these four trials were excluded, and the remaining 476 trials were analyzed.

Performance was quantified using the root-mean-square (RMS) lateral position error of the canoe center, the RMS angular error, and the number of times the canoe hit the river’s edge. Individual and dyadic position error was measured relative to the river centerline, and angular error was measured relative to the local river heading, both using the RMS error. Because the canoe center was fixed vertically on the screen while the river scrolled at *v* = 48 l.u./s, the centerline under S1’s attachment point at the front of the canoe reaches the canoe center only after a delay of Δ = (*L/*2) cos *θ/v*, with *L* = 150 l.u. (Δ ≈ 1.56 s for a straight canoe). The centerline under S2’s attachment point at the rear passed the canoe center Δ earlier. S1’s cursor was therefore compared with the centerline at the canoe center Δ later (lead), and S2’s cursor with the centerline earlier (lag). Time shifts were computed in river time, excluding intervals in which the river was stopped. The power associated with subject *i* is computed as 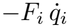 ([22]).

Directional predictive influence between the participants was assessed using conditional Granger causality (GC) between their wrist velocities [23]–To remove influences that both participants received from the river itself, GC was conditioned on the river’s lateral rate and heading rate (the time derivatives of the centerline position and angle). For each trial, intervals in which the river was stopped were removed, and the four signals were low-pass filtered and decimated by a factor of four (to approximately 18 − 20 Hz) and then z-scored. A vector autoregressive model of order 16 (approximately 0.85 s) was fitted to each trial by least squares. GC was computed in both directions, from S1 to S2 and from S2 to S1, in the time domain [24] and in the frequency domain [26].

#### 3) Statistical Analysis

The dyad was the unit of analysis. For the robot-derived measures, trial-level outliers were identified for each measure using the 1.5 × interquartile-range criterion, applied across all trials. A trial that was an outlier on any measure was removed from all of them. Gaze measures were not screened for outliers. The remaining trials were averaged within each dyad, yielding one value per dyad for each combination of the two within-dyad factors: visual view (symmetric, S; asymmetric, A) and haptic feedback (M0– M3). Because performance improved rapidly during the first block of each view condition (Fig. 3a,b), this initial exposure block was excluded from analysis. All comparisons between conditions were thus based on the remaining three blocks of each view. Statistical inference was conducted within a Bayesian framework using the BayesFactor (BF) [27] and *bayestestR* [28] packages in R.

For each measure, we fitted a Bayesian repeated-measures analysis of variance (ANOVA) [29] with visual view and haptic feedback as fixed factors, including their interaction, and dyad as a random factor. Default Jeffreys-Zellner-Siow (JZS) priors were used, with scale parameter *s* = 0.5 for fixed effects and *s* = 1 for the random effect. Bayes factors were estimated using 5 × 10^4^ Monte Carlo iterations. Evidence for each effect is reported as an *inclusion Bayes factor* (BF_incl_), calculated across matched models.

Post-hoc comparisons were conducted using Bayesian paired-samples *t*-tests [30] with the default Cauchy prior on the standardized effect size (scale parameter 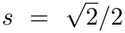). The effect of visual view was assessed by comparing A with S within each haptic feedback mode. The effect of haptic feedback was assessed by comparing every pair of feedback modes separately for each view condition. Each comparison included only dyads with observations in both conditions. Effect sizes are reported as *d*_*z*_ [31], calculated as the mean paired difference divided by the standard deviation of the paired differences.

Time-domain GC was log-transformed and analyzed with a Bayesian mixed model with view, haptic feedback and direction as fixed factors, and dyad and river as random factors. Evidence for each effect is reported as an inclusion Bayes factor. Post hoc Bayesian *t*-tests compared the two directions within each view and compared the directional asymmetry (log GC_*S*1*→S*2_ − log GC_*S*2*→S*1_) between views. GC spectra were compared at each frequency using Bayesian paired *t*-tests on dyad means yielding the *t-test Bayes factor* BF_10_, and only consecutive frequencies with BF_10_ *>* 3 were interpreted.

Bayes factors (BF) were interpreted according to conventional evidence categories [32]: BF_10_ *>* 3 as moderate evidence for a difference, BF_10_ *>* 10 as strong, and BF_10_ *>* 100 as extreme evidence. BF_10_ *<* 1*/*3 was interpreted as moderate evidence for the absence of a difference, whereas values between 1*/*3 and 3 were considered inconclusive.

### C. Computational Model

#### 1) Wrist dynamics

Each participant moves a cursor through the wrist robot, with *c*_*i*_ = −*κq*_*i*_ and *κ* = 250 l.u./rad. With the cursor state **c**_*i*_ = [*c*_*i*_, *ċ*_*i*_]^*⊤*^ and the flexor and extensor activations 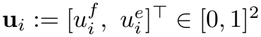, the wrist dynamics are

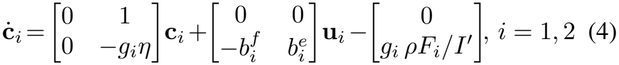

where *I* is the combined human-robot inertia, *I*^*′*^ = *I/κ, η* = *k*_*v*_*/I*, and 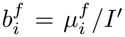 and 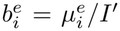, with 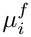 and 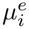 the torque gains of the flexor and the extensor. The last term is the haptic feedback: the coupling force *F*_*i*_ of Eq. 2, rendered as a wrist torque only when *g*_*i*_ = 1. It depends on the canoe through the attachment point *x*_*i*_ of Eq. 1 and its velocity 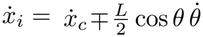. Here and below, the upper sign refers to S1 and the lower sign to S2. The resulting cursor positions change the state of the canoe governed by Eq. 3.

#### 2) Clearance from the river edges

The river is described by its centerline *x*_*r*_, its right and left edges *x*_*R*_ and *x*_*L*_, and its local tangent angle with the vertical *ψ*, all functions of the river time *t*_*r*_, which advances with time *t* while the river moves and stops while it is stopped. The river point at the canoe center at river time *t*_*r*_ is *x*_*r*_(*t*_*r*_); the point beside each attachment point is *x*_*r*_(*t*_*r*_ ± Δ), with the lead Δ = (*L/*2) cos *θ/v* of Section II-B, and the centerline moves laterally at the rate 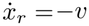 tan *ψ*. In the frame of the canoe, the edges at a river-time offset *d* from the canoe center lie at

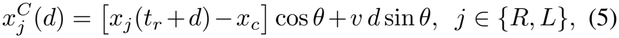

and the clearance of the canoe from the edges is

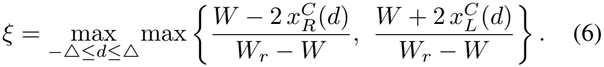

The clearance is *ξ* = − 1 when the canoe lies on the centerline and along the river, *ξ* = 0 when it touches an edge, and *ξ >* 0 when it crosses one. In the model, the maximum was taken over 21 points along the canoe.

#### 3) Control

Each participant is modeled by a two-layer controller. An outer layer sets the commanded cursor velocity 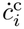 from what the participant sees, and an inner layer drives the wrist muscles so that the cursor follows this velocity. The two controllers act independently, and each participant influences the other only through the canoe. Given our results on the limited influence of the haptic feedback (see Section III), the model focuses on simulating the effects of the visual information conditions.

##### a) Outer layer: commanded cursor velocity

Each participant observes the canoe and the river with a delay *t*_*d*_. Writing 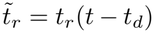 and 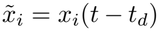 for the river time and the attachment point seen at time *t*, the commanded velocity is

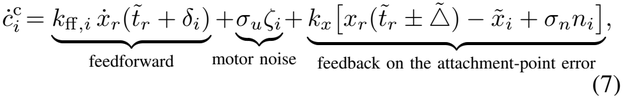

where 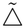 is the lead for the canoe angle seen at *t* − *t*_*d*_. The feedforward term follows the river’s lateral rate at a preview *δ*_*i*_ from the canoe center. The feedback term moves the cursor so as to bring the participant’s own attachment point onto the centerline beside it. Because the wrist integrates the commanded velocity, this acts as integral control of the attachment-point error. To first order, this error combines the position and heading errors of the canoe,

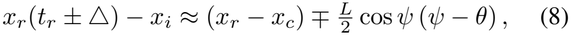

so no separate heading term is needed. The observation noise *n*_*i*_ is a first-order Gauss-Markov process with unit variance and correlation time *t*_*n*_, and *ζ*_*i*_ ∼ *N* (0, 1) is white motor noise drawn at every controller update.

S1 and S2 share the delay *t*_*d*_, the feedback gain *k*_*x*_ and the noise levels *σ*_*n*_ and *σ*_*u*_; they differ only in the feedforward term. S1 follows the river’s rate with unit gain at its own attachment point (*k*_ff,1_ = 1, *δ*_1_ = *L/*2*v* = 1.56 s), whereas S2 has its own gain *k*_ff,2_ and a view-dependent preview *δ*_2_. Under asymmetric view, S2 could not see the river ahead of the canoe center, so for *δ*_2_ *>* 0 only the slope at the center would be available to S2. The fitted *δ*_2_ was negative under both views (Table II).

**TABLE II:**
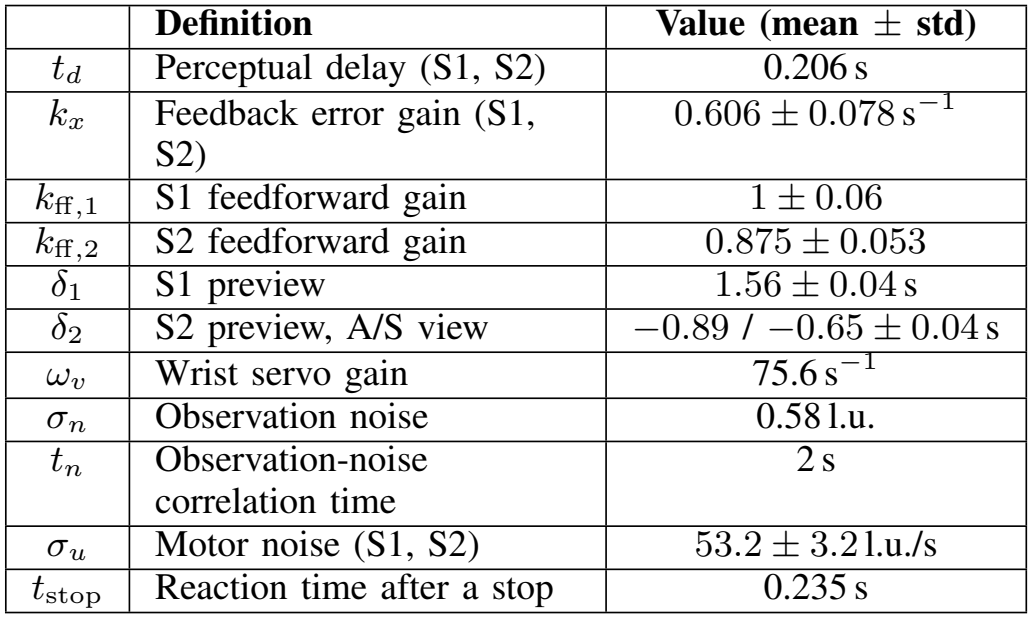
Model parameters. Gains and motor noise vary lognormally, and their standard deviation is given to first order.

|  | **Definition** | **Value (mean ± std)** |
|---|---|---|
| *t<sub>d</sub>* | Perceptual delay (S1, S2) | 0.206 s |
| *k<sub>x</sub>* | Feedback error gain (S1, S2) | 0.606 ± 0.078 s<sup>−1</sup> |
| *k*<sub>ff,1</sub> | S1 feedforward gain | 1 ± 0.06 |
| *k*<sub>ff,2</sub> | S2 feedforward gain | 0.875 ± 0.053 |
| *δ*<sub>1</sub> | S1 preview | 1.56 ± 0.04 s |
| *δ*<sub>2</sub> | S2 preview, A/S view | −0.89 / −0.65 ± 0.04 s |
| *ω<sub>v</sub>* | Wrist servo gain | 75.6 s<sup>−1</sup> |
| *σ<sub>n</sub>* | Observation noise | 0.58 l.u. |
| *t<sub>n</sub>* | Observation-noise correlation time | 2 s |
| *σ<sub>u</sub>* | Motor noise (S1, S2) | 53.2 ± 3.2 l.u./s |
| *t*<sub>stop</sub> | Reaction time after a stop | 0.235 s |

##### b) Inner layer: wrist velocity servo

The muscles drive the cursor velocity towards 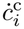 and cancel the haptic force:

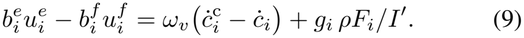

Only one muscle is active at a time: with *a*_*i*_ the right-hand side of Eq. 9, 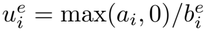 and 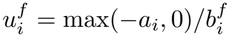^*f*^, both saturated to [0, 1]. Substituting Eq. 9 into Eq. 4 gives a first-order velocity servo,

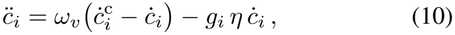

with a time constant 1*/ω*_*v*_ ≈ 13 ms, much faster than the outer layer.

##### c) River stops

As in the experiment, the river stops when the canoe touches an edge. Using the clearance of Eq. 6 for the canoe’s long axis (*W* = 0), the river stops when *ξ* ≥ − 0.07 and restarts when *ξ <* − 0.26, the median values at the recorded stops. While the river is stopped, each participant waits for a reaction time *t*_stop_ and then moves their attachment point to where it would be if the canoe were centered on, and aligned with, the river, using the feedback gain of Eq. 7 on the current state:

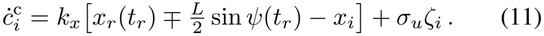

##### d) Implementation and fitting

The canoe and wrist dynamics were integrated with a fourth-order Runge-Kutta scheme with a 2.5 ms step, and the outer layer was updated every 12.5 ms with a zero-order hold. Each virtual dyad ran the same trials as the corresponding real dyad, with its own small variation of the gains, previews and motor noise. The gains, times and noise levels in Table II were identified by matching the virtual dyads to the real ones in blocks 2–4, using six metrics: the canoe’s position and angular errors, and the cursor errors and force magnitudes of S1 and S2. For each metric, the difference between the model and human means was divided by the standard error of the human mean, giving a mismatch *z*, and the cost was

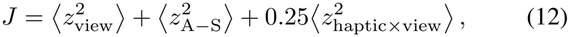

Where ⟨·⟩ is the average over the metrics of the squared mismatch of the two view means, of the difference between the views, and of the eight view × haptic mode cells, respectively, the latter down-weighted because each holds fewer dyads. Thus *J* ≈ 1 means that the model lies within the uncertainty of the human data. *J* was minimized by a stochastic search over the parameters, and the variation of the parameters across dyads was then fitted so that the spread of the virtual dyads matched that of the real ones.

## III. Results

Fig. 2 illustrates a river bend together with the trajectories of the canoe and both cursors. Grouping the sessions by the order of the symmetric and asymmetric blocks shows that adaptation under the asymmetric view transferred to the symmetric view (Fig. 3b), but not the other way round (Fig. 3a). Moreover, participants adapted the task relatively quickly during the exposure block of each view (block 1 or 5), and performance did not change significantly in the remaining blocks.

**Fig. 2:**
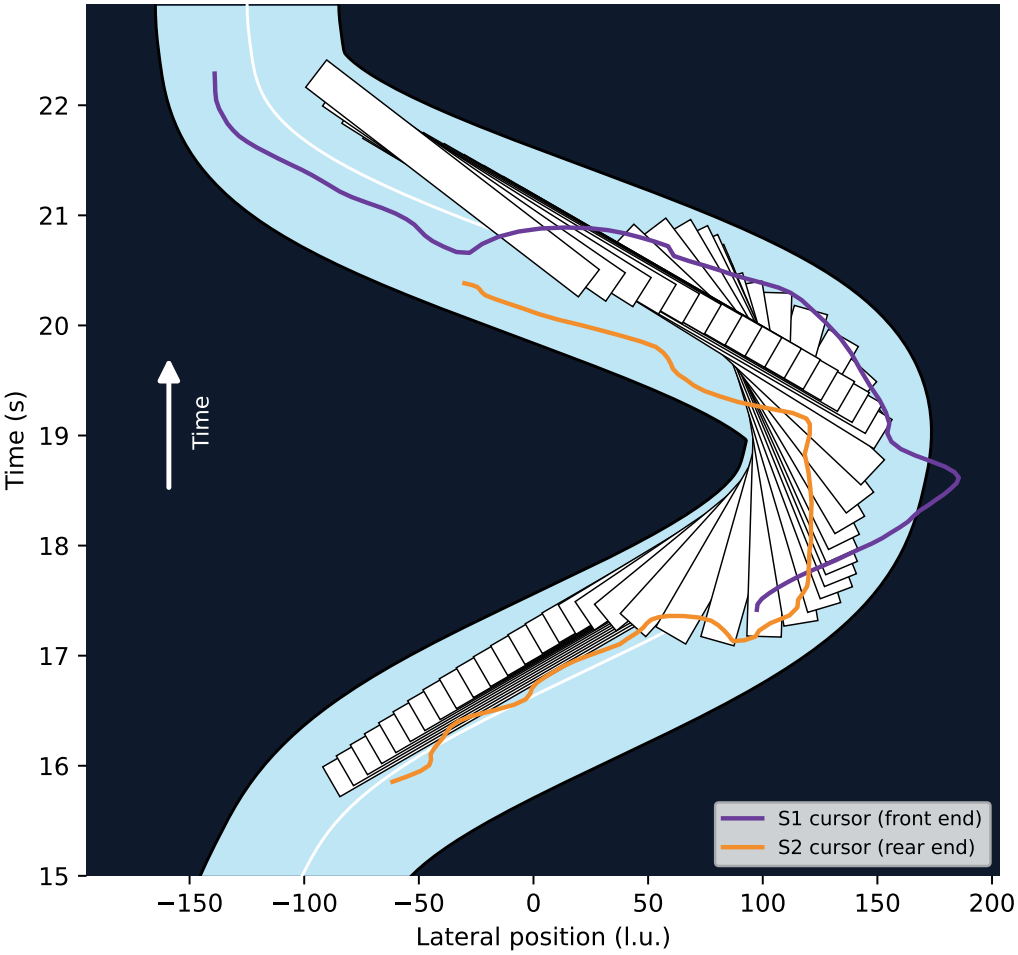
Representative of the virtual task. A portion of a trial (dyad 3, asymmetric view, M3). The canoe is shown every 0.1 s, and the cursor trajectories are sampled every 0.05 s. In the river reference frame, the canoe advances upward with time (in the experiment, the canoe is fixed vertically and the river descends).

**Fig. 3:**
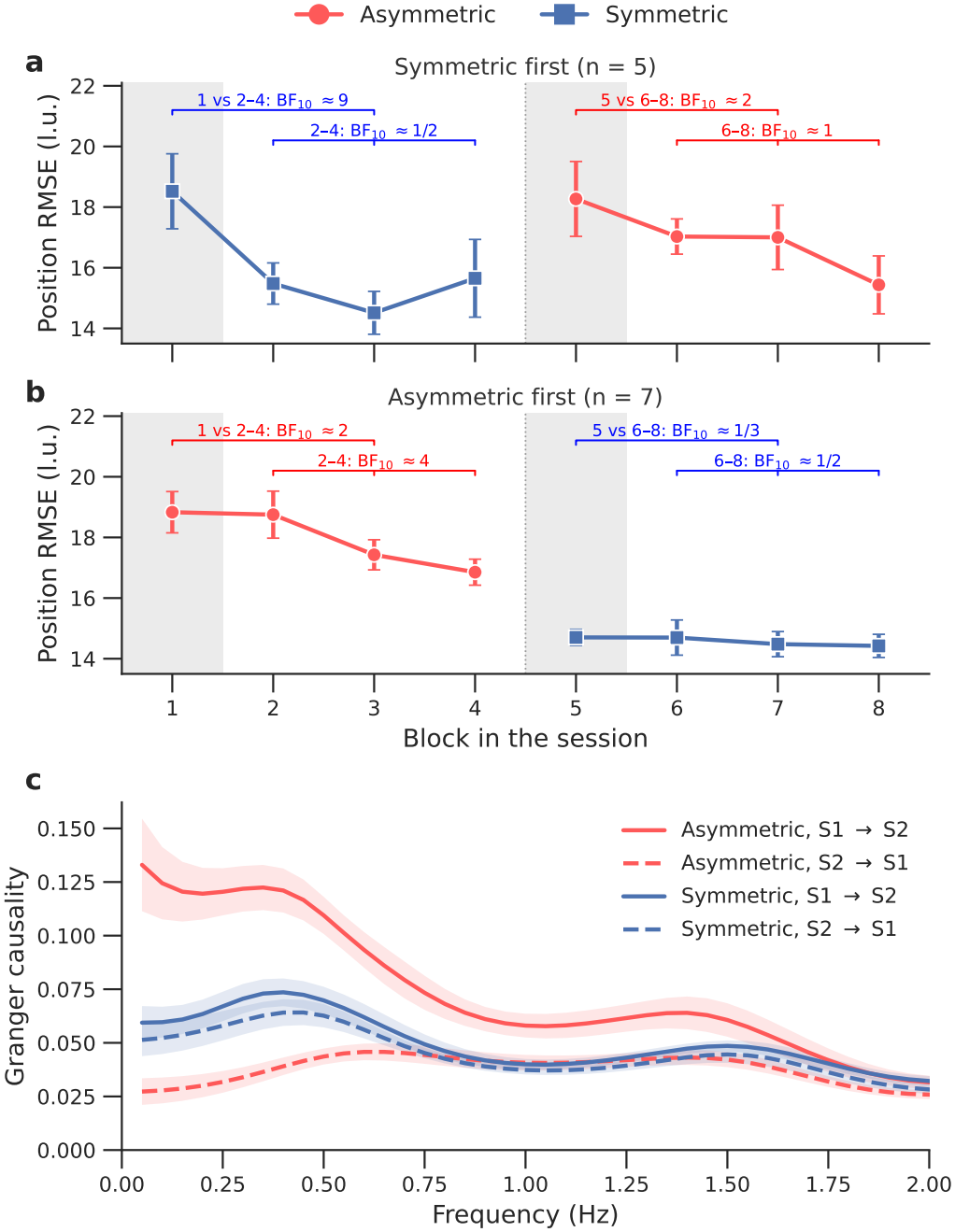
Learning and directional influence. **(a,b)** Lateral position RMSE of the canoe w.r.t. river centerline across the eight blocks of the session, for dyads that started with symmetric view (a) and with asymmetric view (b). Grey areas mark the first block of each view, which is excluded from the analysis of Fig. 4. **(c)** Conditional spectral Granger causality between the wrist velocities of S1 and S2, conditioned on the river and pooled over haptic modes and all blocks. Solid lines show S1 to S2 and dashed lines S2 to S1. Markers and lines show the mean across dyads, and error bars and shaded bands show *±*1 standard error.

*Conditional Granger causality* (GC) between the participants’ wrist velocities was stronger from S1 to S2 than from S2 to S1 under asymmetric view (BF_10_ ≈ 151, *d*_*z*_ = 1.56; Fig. 3c). This indicates that the front partner more strongly influenced the rear partner than conversely. On the other hand, the evidence was inconclusive under symmetric view (BF_10_ ≈ 1*/*2, *d*_*z*_ = 0.36). This suggests that S1 took a leading role when only S1 could see the upcoming river but S2 not. Because S1 sat at the front of the canoe in both view conditions, the leadership arose from the difference of information available to the two participants rather than from their positions within the canoe.

### Overall performance

Despite the more demanding asymmetric view, the dyads performed the task consistently under both views and all haptic feedback modes. The number of times the canoe hit the river edge did not differ between asymmetric and symmetric view (BF_incl_ ≈ 1*/*4, *d*_*z*_ = 0.04) or across haptic modes (BF_incl_ ≈ 1*/*8; Fig. 4a).

**Fig. 4:**
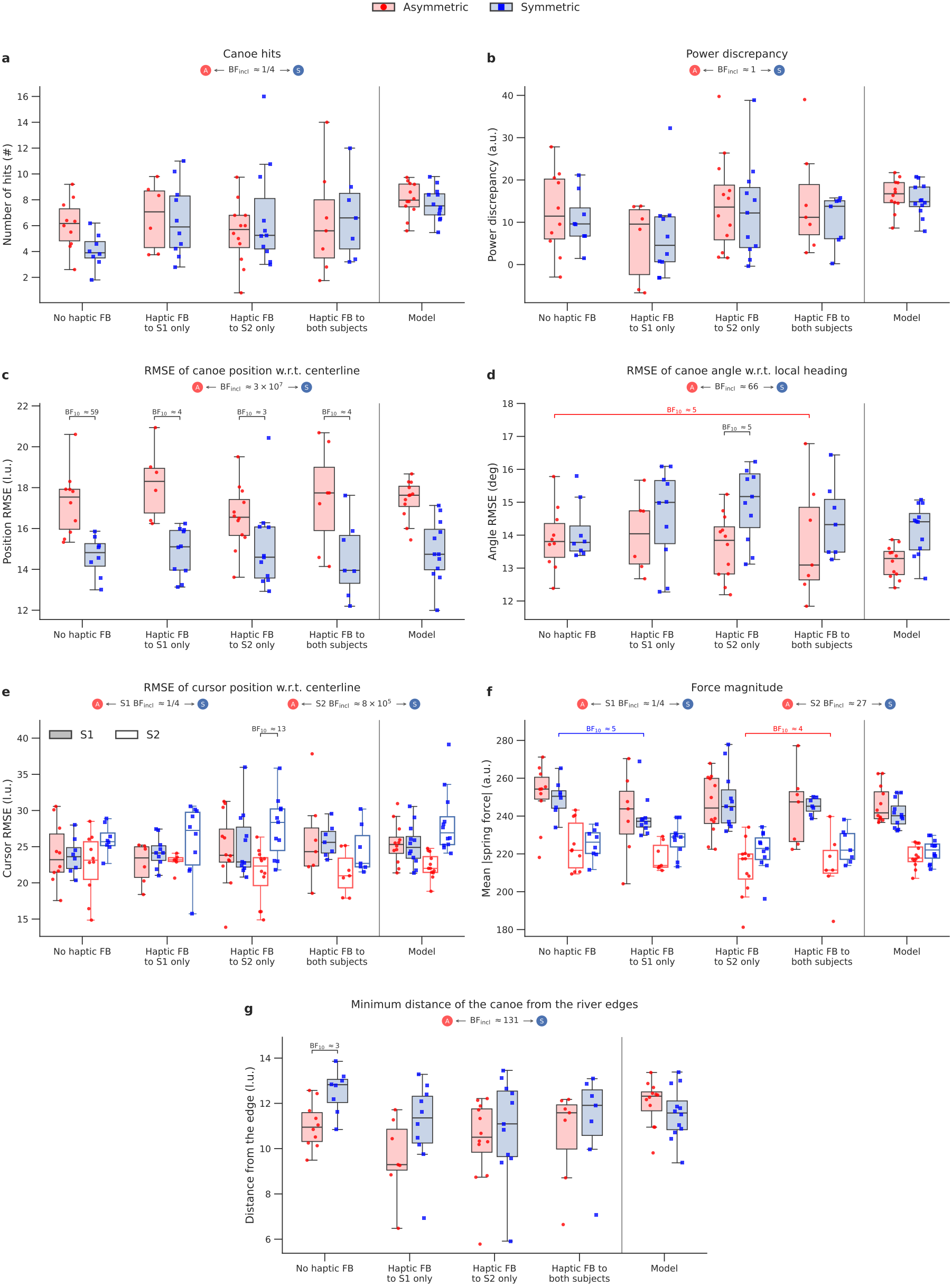
Performance and strategies across view and haptic feedback conditions. Each panel shows one measure for the four haptic feedback modes under asymmetric (red) and symmetric (blue) view. **(a)** Number of times the canoe hit the river edge. **(b)** Power discrepancy between the participants. **(c)** RMSE distance of the canoe’s lateral position with respect to the river centerline. **(d)** RMSE distance of the canoe’s angle with respect to the local river heading. **(e)** RMSE distance of each participant’s cursor position with respect to the river centerline at that participant’s attachment point. **(f)** Mean magnitude of the spring force of each participant’s coupling. **(g)** Distance between the canoe and the nearest river edge, averaged over trials after excluding the exposure block of each view. The badge beneath each title gives the inclusion Bayes factor for the effect of view.

Although S1 became the leader under asymmetric view, the evidence on whether the power discrepancy between the participants changed with view was inconclusive (BF_incl_ ≈ 1, *d*_*z*_ = 0.47; Fig. 4b). The emergence of the leader-follower roles therefore did not come with a clear shift in the balance of power between the partners.

### Task-space strategy

Under symmetric view, when both participants could see the upcoming river, the dyads kept the canoe closer to the river centerline: the position error was smaller than under asymmetric view (BF_incl_ ≈ 3 × 10^7^, *d*_*z*_ = 1.73), in all four haptic modes (BF_10_ ≈ 59, 4, 3, 4 for M0–M3; Fig. 4c).

Under asymmetric view, dyads appeared to rely less on angular displacements, which may be a sign that asymmetric knowledge induced a lower coordination to rotate the canoe in turns. Indeed, angular error relative to the local heading was smaller in asymmetric than symmetric (BF_incl_ ≈ 66, *d*_*z*_ = 1.24; Fig. 4d). The view effect on angular error depended on the haptic mode (BF_incl_ ≈ 3) and was clearest with feedback to S2 only (BF_10_ ≈ 5, *d*_*z*_ = 0.92). Haptic feedback had no overall effect on angular error (BF_incl_ ≈ 1*/*9) or position error (BF_incl_ ≈ 1*/*7).

### Participants’ control strategies

S1’s strategy changed little across views and haptic modes, whereas S2’s was affected by the view (Fig. 4e). S1’s cursor error did not differ between views (BF_incl_ ≈ 1*/*4, *d*_*z*_ = 0.04) or haptic modes (BF_incl_ ≈ 1*/*7), while S2’s cursor error was smaller under asymmetric view (BF_incl_ ≈ 8 × 10^5^, *d*_*z*_ = 2.20). This is consistent with S1 planning ahead in the same way under both views while S2 adapted to S1’s inputs with a delay (as also shown with Granger causality), which interaction can be classified as assistance cooperation [10].

S1 took the lead and exerted more effort, with a larger force magnitude than S2 under both views (asymmetric: BF_10_ ≈ 208, *d*_*z*_ = 1.63; symmetric: BF_10_ ≈ 102, *d*_*z*_ = 1.48; Fig. 4f). S1’s force did not differ between views (BF_incl_ ≈ 1*/*4, *d*_*z*_ = 0.00), consistent with a view-independent strategy. Overall, S2 exerted less force than S1, and they further reduced their input force under asymmetric view (BF_incl_ ≈ 27, *d*_*z*_ = 0.62), although this difference was not consistent across dyads.

Finally, we quantified how close the canoe came to the river’s edges, which can be seen as the safety margins the participants adopted either volitionally or due to the visual feedback constraints (Fig. 4g). Under asymmetric view, the canoe moved closer to the edges than under symmetric view (BF_incl_ ≈ 131, *d*_*z*_ = 0.86). This difference was clearest without haptic feedback (BF_10_ ≈ 3, *d*_*z*_ = 1.08).

### Gaze

Figure 5a shows the distribution of S1’s gaze on the monitor in the two views. The lateral (x) distributions were nearly identical in the two views as the lateral gaze position did not differ between the views (BF_incl_ ≈ 1*/*4, *d*_*z*_ = 0.02). The vertical (y) distributions exhibited a major peak ahead of the canoe. In the asymmetric view, the whole longitudinal distribution was shifted towards the upcoming river compared to the symmetric view (BF_incl_ ≈ 6 × 10^5^, *d*_*z*_ = 0.98), which suggests a longer planning horizon in asymmetric settings. Moreover, S1 made more frequent saccades under asymmetric than symmetric view (BF_incl_ ≈ 40, *d*_*z*_ = 0.56; Fig. 5b), although no single haptic mode showed a clear difference. Saccade amplitude was smaller under asymmetric view (BF_incl_ ≈ 66, *d*_*z*_ = 0.80; Fig. 5c), most clearly with feedback to S2 only (BF_10_ ≈ 6, *d*_*z*_ = 1.04).

**Fig. 5:**
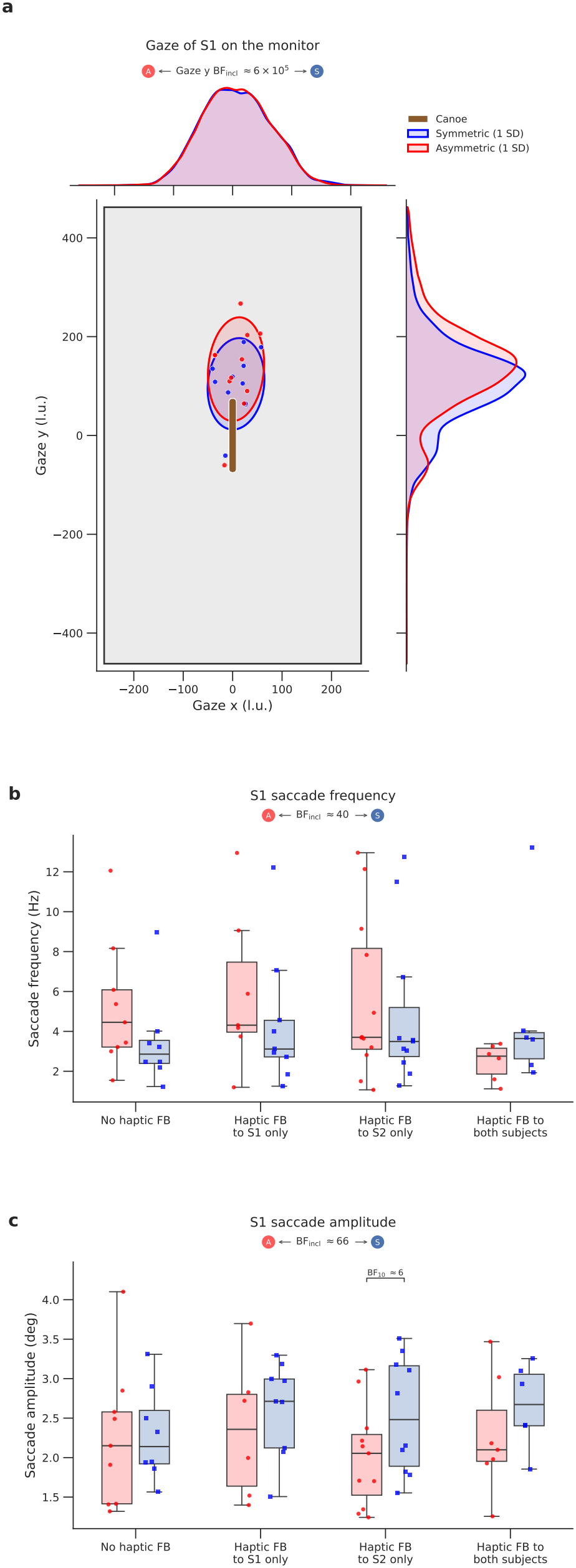
S1’s average gaze location on the monitor, saccade frequency and saccade amplitude. **(a)** Gaze of the front participant (S1) on the monitor (520 × 924 l.u.), with the canoe drawn to scale at the screen center. Ellipses show one standard deviation of the gaze samples in the symmetric (blue) and asymmetric (red) views, with each dyad weighted equally; dots show each dyad’s mean gaze. The top and right panels show the distributions of the lateral (x) and longitudinal (y) gaze positions. Gaze was directed further ahead along the river in the asymmetric view. One dyad was excluded because of a calibration offset. **(b**,**c)** S1’s saccade frequency and amplitude.

## IV. Discussion

Despite partial and asymmetric visual information, humans can seamlessly co-manipulate large objects in cluttered environments, such as moving a piano through a doorway. We therefore expected participants to perform the co-manipulation task successfully. In our experiment, dyads controlled a relatively long object governed by physical dynamics while navigating through a randomly bending river. Importantly, one condition restricted visual feedback to mimic real-world carrying tasks, in which one partner can see the path ahead while the other can monitor the space around the object. The low number of boundary crossings even in the early trials indicates that (i) participants successfully performed the task, and (ii) learned it relatively quickly, which both support our first hypothesis (H1).

Conditional Granger causality (GC) was used to analyze the direction of influence between the partners’ movements (Fig. 3c), following the approach proposed in [25]. Whereas earlier metrics based on the power discrepancy between partners [22] showed no clear difference, the GC analysis indicated that the participant with more task relevant visual information assumed a leading role. This finding supports H3, which predicted that leadership would emerge from information asymmetry. Previous studies have shown that leader-follower roles can emerge when mechanical constraints differ between partners [9], [15], when they have different levels of sensory acuity [16], [17], or when only one of them is made aware of the task’s goal [18]. Our results extend these findings by showing that leadership is context dependent and is determined by the task relevance of available sensory information.

Interestingly, participants transferred what they had learned under restricted, complementary vision (asymmetric view) to the full shared vision (symmetric view), but not in the opposite direction (Fig. 3a,b). This suggests that the motor skills required under symmetric visual condition are contained within those required under asymmetric condition. In other words, successful performance in asymmetric settings requires additional coordination strategies that are not necessary when both partners can see the upcoming river.

Participants also adopted different control strategies depending on the information available to them. With full visual information, they appeared to continuously update their actions based on the observed river and adjust their cursor movements to keep the canoe close to the centerline (Fig. 4c-e). This behavior suggests that they may have performed the task by focusing on environmental constraints rather than inferring their partner’s actions, thereby reducing the associated cognitive demands [33]. Under restricted vision, the rear participant may have played a more prominent role by compensating for the actions of the front participant. This interpretation would be consistent with the lower angular RMSE observed under restricted than under full vision.

The front participant also adapted their interpretation of the environment to the information available to them. Under asymmetric viewing, when only the front participant could see the upcoming river, they directed their gaze further ahead along the river than in the symmetric view (Fig. 5a) and maintained it there more steadily, with more frequent and smaller saccades (Fig. 5b,c). This pattern is consistent with the front participant using a longer time horizon to plan their contribution to the canoe’s manipulation. As shown in Fig. 4g, the canoe moves closer to the riverbank in the asymmetric condition, suggesting a reduced safety margin. Two possible explanations emerge from this observation. First, the back participant may have focused on correcting the front participant’s rotations while fully exploiting the river’s width. This would allow the front participant to look farther ahead, entrusting their partner with maintaining a safe trajectory. Second, the back participant may have been unable to contribute accurately because they lacked information about upcoming environmental constraints. This interpretation is supported by the back participant’s overall lower contribution in the asymmetric condition, both in terms of cursor movements and the force applied on the canoe, as well as by their lower overall angular RMSE. It also explains why they adopted a follower role. Consequently, their input became less influential in the task, prompting the front participant to plan farther ahead either to refine their predictions of the back participant’s actions or to allow more time to prepare for upcoming river turns. Together, these findings support H2, which predicted that access to taskrelevant information would affect both control and observation strategies.

Providing haptic feedback did not substantially affect task performance. One possible explanation is that, although the visual information was restricted in the asymmetric condition, it was sufficiently precise for participants to control the canoe without relying strongly on haptic feedback. Future experiments could test this explanation by degrading visual information, for example by blurring the canoe or the river. Another possibility is that participants used an internal model of the viscoelastic coupling between each cursor and the canoe. Because both cursors and the canoe remained visible in all conditions, participants could potentially predict the interaction forces from visual information alone, rendering haptic feedback largely redundant. Thus, H4 may not hold when visual information is sufficiently precise.

These observations suggest a critical role of (i) the planning ability of the two partners, and (ii) visual feedback delays, as they induce a delay in the ability of S2 to adapt their motion, forcing them to adopt a low-activity follower role. As such, we propose a model focusing on these two parameters, without accounting for high-level cognitive abilities, which could include internal models of the task’s geometry and complex inference of the partner’s behavior. In this model, each partner commands the velocity of their cursor based on visual information delayed by approximately 0.2 s, which corresponds to reaction times previously observed in similar settings [7]. The commanded velocity combines the river’s lateral velocity at a preview point with a feedback correction based on the discrepancy between the river centerline and the participant’s end of the canoe. The wrist muscles then track this command through a fast velocity servo. Both partners have the same delay and feedback gain and differ only in their visual preview: S1 looks ahead of the canoe and S2 behind the canoe center. Despite its simplicity, the model reproduced the main characteristics of the human data, including the canoe’s positional and angular errors, the partners’ cursor errors and force magnitudes, and the differences between the asymmetric and symmetric visual conditions (Fig. 4). Specifically, 75% of our low-level model’s condition means fell within two standard errors of the corresponding human means, and the difference between visual conditions was captured by changing S2’s preview alone. However, the model did not reproduce the difference between the symmetric and asymmetric conditions in terms of safety margins (Fig. 4g). Predicting this difference would require to refine the model by including the whole task geometry and optimal motion planning. This extension was beyond the scope of the present paper, where we focused on the impact of the information available to partners and sensorimotor delays. Our results suggest that during object co-manipulation, humans’ movement patterns are largely impacted by these two characteristics, possibly with a lower impact of the general movement strategy itself (in the sense of the costs minimized by the central nervous system). Note that these observations may be impacted in tasks that may be slower, more effortful, or if collisions are not permitted because too hazardous.

## V. Conclusion

This study examined how two partners coordinate the manipulation of a large object within a constrained environment when they receive restricted but complementary visual information about the object’s front and rear, with haptic feedback provided to either or both partners. Dyads performed the task reliably across all conditions, demonstrating that partners can compensate for incomplete information about the environment. When only the front partner could see the path ahead, conditional Granger causality revealed a stronger influence from the front, informed partner; when both partners could see ahead, influence was more balanced. The lessinformed partner did not become passive but reduced their overall contribution to the task to allow a safe execution by facilitating the task of the informed partner. As a consequence, the front partner shifted their gaze further ahead along the path and held it there more steadily than when provided full view, taking on the role of anticipating the path for the dyad. Haptic feedback had little effect on performance, possibly because the mechanical coupling between the cursor and canoe could be inferred from visual information. These findings show that leader-follower roles can emerge from information asymmetry during co-manipulation of a large object in a constrained environment, and that the information available to each partner shapes both how the dyad controls the object and how each partner observes the task. These results can inform the design of sensory-feedback strategies for human-robot cooperative systems, in which partners often have access to different views of the task.

## Author Contributions

A.K.: Conceptualization, methodology, formal analysis, visualization, computational modeling, supervision, writing.

F.H.: Conceptualization, methodology, unity development, investigation, data acquisition, visualization, initial modeling, writing.

D.V.: Conceptualization, writing.

E.B.: Conceptualization, supervision, writing.

## Acknowledgments

We thank all participants in the study, and Yihan Liu for helpful suggestions on the experiment.

